# Overcoming non-small cell lung cancer radiation resistance by modulating the tumor-immune ecosystem

**DOI:** 10.1101/458372

**Authors:** S. Nizzero, J.C.L. Alfonso, A. Álvarez-Arenas, I. Mirzaev, I.K. Zervantonakis, T. Lewin, A. Rishi, E. Piretto, T.V. Joshi, D.N. Santiago, A. Karolak, R. Walker, F.A. Karreth, J. Torres-Roca, H. Enderling

## Abstract

Non-small-cell lung cancer is the leading cause of cancer death worldwide. Although radiotherapy is an effective treatment choice for early-stage cases, the 5-year survival rate of patients diagnosed in late-stages remains poor. Increasing evidence suggests that the local and systemic effects of radiotherapy dependent on the induced anti-tumor immune responses. We believe that an educated adaptation of radiotherapy plans based not only on the induced immune responses, but also on the tumor-immune ecosystem composition at the beginning of treatment might increase local tumor control. We propose two different mathematical models to evaluate the potential of the tumor-immune context to inform adaptation of treatment plans with the aim of improving outcomes.

## I. Background

Lung cancer is the second most common cancer, with an estimated of 224,390 new cases (117,920 in men and 106,470 in women) and 158,000 deaths in the US in 2016 [1]. About 24% of patients present with Stage III non-small cell lung cancer (NSCLC) for which chemoradiotherapy is the standard-of-care for inoperable cases. A systems biology model of tumor intrinsic radiosensitivity, radiosensitivity index RSI [2], has been clinically evaluated in over 8,000 patients in multiple disease sites, including lung. Whilst 60% of stage III NSCLC patients with lower RSI values (RSI < 0.31) are controlled by radiation, treatment-resistant patients fail locoregionally with a 5-year overall survival of 5-14%. Patients with relapsed NSCLC are frequently reirradiated, rarely with improved outcomes. This offers an opportunity for investigating alternative radiation approaches, either alone or in combination with targeted agents.

Multiple altered radiation fractionation protocols, including the accelerated fractionation or hyperfractionation have been tested clinically without significant outcome improvements, in part due to (i) non-specific selection of patients into the different treatment protocols, and (ii) the prevailing dogma of delivering the same dose per fraction and constant fractionation intervals. Combination of radiation and targeted therapy, either concurrently or sequentially, has achieved significantly prolonged overall survival in patients with confirmed mutation status [3]. Compelling results are also emerging from clinical trials evaluating combination of radiotherapy with immunotherapy [3]. Interestingly, radiation therapy also induces cell stress and immunogenic cell death, thereby exposing a wealth of previously hidden tumor-associated antigens, heat shock or stress proteins (HSP) and danger-associated molecular patterns (DAMP), which are endogenous immune adjuvants that can elicit an antitumor immune response [4]. To date, radiation therapy and dose scheduling has not specifically focused on enhancing the immune response to tumor antigens. Here we propose a novel investigation in which we seek to exploit the immunostimulatory consequence of radiation therapy to induce de novo antitumor immune responses and augment the effect of blocking immunosuppression.

We hypothesize that the tumor-immune ecosystem composition at clinical presentation and its evolution during treatment contributes to clinical response to radiotherapy. Radiation-induced promotion of adaptive immunity, either stimulation of antitumor immunity or inhibition of immune regulatory mechanisms, may be required to overcome radiation resistance, particularly in Stage III NSCLC patients. It is conceivable that alternative radiation fractionation, either alone or in combination with targeted therapy, may better synergize with the host immune system to control tumors resistant to conventional protocols.

## II. Material And Methods

Transformed cancer cells are confronted with an innate and adaptive immune surveillance. Tumor-associated antigens (HSPs and DAMPs) released during cell death are endogenous immune adjuvants that can both initiate and continually stimulate an immune response against tumors. In retaliation, tumors can hijack intrinsic immune regulatory programs that are intended to prevent autoimmune disease, thereby facilitating continued growth despite the activated antitumor immune response. Irradiation induces cell stress and immunogenic cell death, thereby exposing a wealth of previously hidden and new tumor associated antigens to the immune system. However, radiation also induces cell death in the tumor inhibiting and immune inhibiting immune populations, which makes predicting radiation response a complex dynamical system that could be best understood with the help of mathematical modeling (Figure 1). To that end, we develop an agent-based model (ABM) to account for the spatial aspects of tumor growth and immune cell dynamics, as well as formulate an ODE model with the potential to scale up to realistic cell population sizes.

**Fig. 1.**
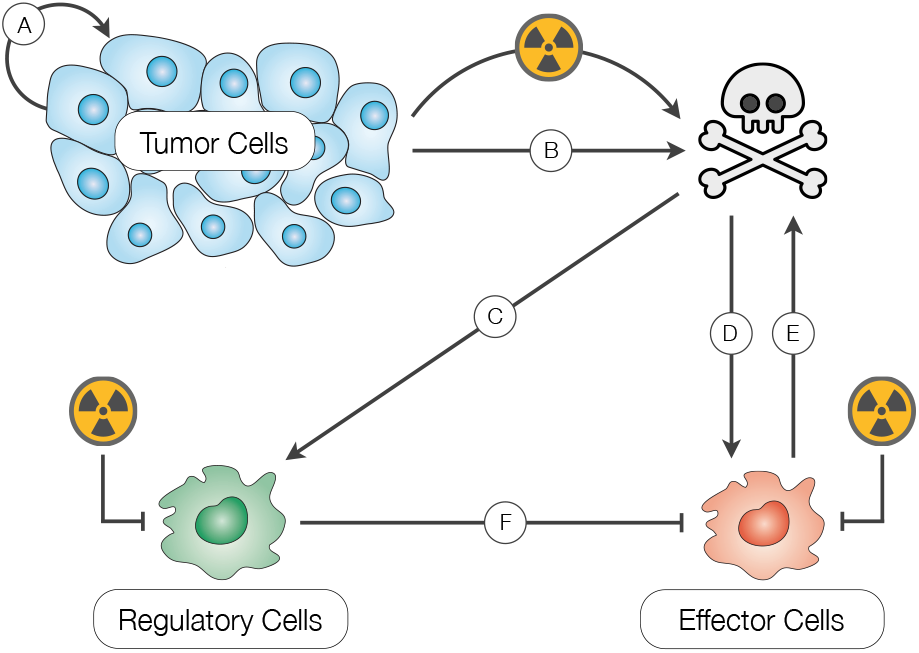
Diagram of the system interactions between tumor, immune effector and immune regulatory and the response to radiation therapy.

### ODE model

We define the system variables as cancer (*T*), immune effector (*E*) and immune regulatory (*R*) cells, and their respective dynamics with

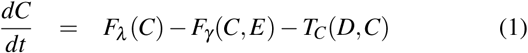

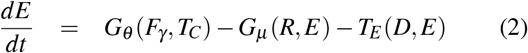

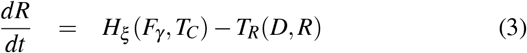

where the functions *F_λ_* represents the net tumor growth, *F_γ_*(*C*, *E*) is the death of tumor cells due to the cytotoxic action of effector cells, and *G_μ_* (*R*, *E*) represents the inactivation of effectors due to regulatory cells. The functions *G_θ_* (*F_γ_*, *T_C_*) and *Hξ* (*Fγ*, *T_C_*) are the recruitment of effector and regulatory cells induced by radiation-induced immunological tumor cell death, respectively. The treatment-associated functions *F_T_* (*D*, *T*), *F_E_* (*D*, *E*) and *F_R_* (*D*, *R*) represent the death fraction of each cell population irradiated with an acute dose *D*.

### ABM model

To capture spatial interactions between cells, we simulate the tumor-immune ecosystem under radiotherapy using a lattice-based agent-based model (ABM). Cells are individually represented and its fates determined by mechanistic rules. Each cell in the model is capable of proliferate. Moreover, immune effector cells are capable of killing tumor cells by direct contact. Similarly, immune regulatory cells can kill effector cells by direct contact. The ABM model also accounts for the RSI values corresponding to lung cancer [2], as well as the recruitment of effector and regulatory cells to the tumor region.

## III. Results

Preliminary Results of the ODE and ABM models indicate that the regulatory to effector cell ratio (Treg/Teff) has a significant impact on the success of radiation therapy. Figure 2 and 3 show the tumor and immune cell evolution before and after radiotherapy for different recruitment rates of regulatory cells. We found that increasing values of Treg/Teff adversely impact treatment outcomes by reducing the effectiveness of radiation therapy, i.e. the tumor microenvironment exhibits stronger suppressive activity mediated by regulatory cells. These *in silico* findings suggest that a personalized treatment adaptation based on the pre-treatment tumor-immune ecosystem and its evolution during radiation therapy might result in better outcomes.

**Fig. 2.**
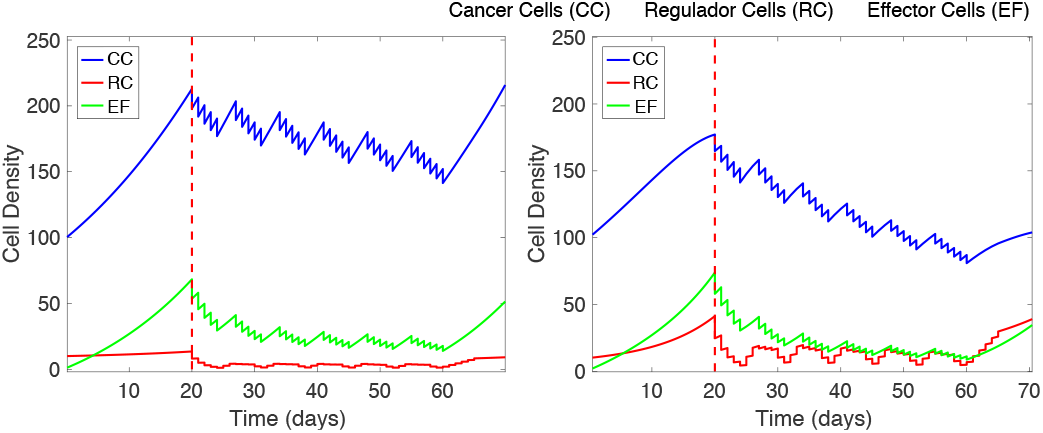
ODE model for tumor-immune dynamics under radiotherapy. Results from simulations with a low (*Left*) and high (*right*) recruitment rate of regulatory cells.

**Fig. 3.**
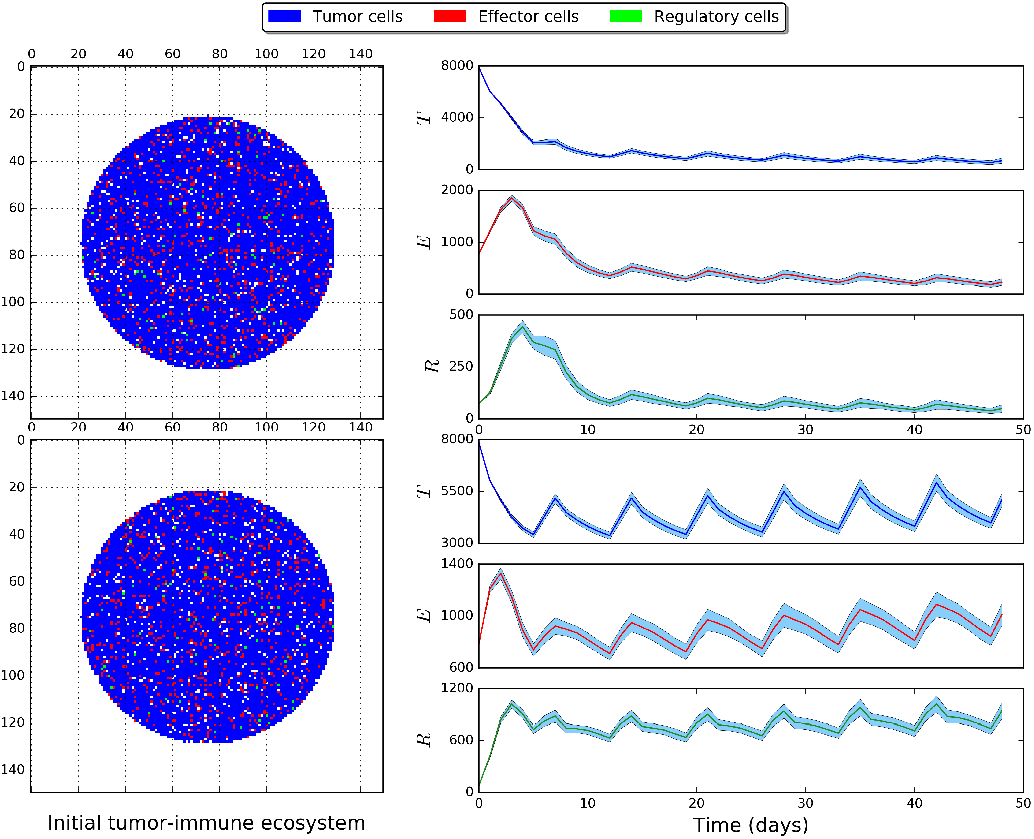
Lattice-based ABM model for tumor-immune dynamics under radiotherapy. Results from simulations with a low (*TOP*) and high (*BOTTOM*) recruitment rate of regulatory cells. Filled region represents 95% confidence interval for 100 different simulations for the same initial conditions.

## IV. Discussion & Future Work

Preliminary model simulations provide a proof-of-concept demonstration of the importance of tumor-immune ecosystem dynamics in determining clinical response to radiotherapy. This lays the foundation for investigations into adaptive therapy to overcome radiation resistance in patients who failed conventional treatment. Whilst future translation of project findings into the clinic would most likely be for the subgroup of locally recurrent Stage III NSCLC patients subject to re-irradiation, we anticipate this project to also provide novel insights into combining RSI and immune infiltration evaluation to identify individual patients who are likely to fail conventional therapy, and thus are likely candidates for immediate treatment adaptation.

## V. Acknowledgments

We would like to thank the IMO Chair, Dr. Alexander Anderson, for organizing the 6th Annual Moffitt IMO workshop: Resistance, where this project was conceived. We are also extremely grateful to the Moffitt Cancer Center and the Moffitt PSOC for supporting this workshop through the NCI U54CA193489 grant.

## Notes

This research was supported by H. Lee Moffitt Cancer Center & Research Institute, Tampa, FL 33612 USA

